# Bridging Hearts and Muscles: Genetic Insights into Cardiac Structure and Sarcopenia

**DOI:** 10.1101/2025.04.29.651359

**Authors:** Chao Wang, Wenqing Xie, Yong Zhu, Zhi Liu, Guang Yang, Da Zhong, Yusheng Li, Shushan Zhao

**Author notes:** These authors contributed equally (Chao Wang, Wenqing Xie). **Correspondence to:** Shushan Zhao; Yusheng Li **Shushan Zhao** **Yusheng Li**.

## Abstract

**Background:** Currently, the link between cardiovascular diseases and sarcopenia is increasingly garnering attention from researchers. However, studies exploring the association between cardiac structure and sarcopenia are sparse. This study aims to investigate the potential genetic links between cardiac structural features and sarcopenia.

**Methods:** During the discovery phase, we employed Linkage Disequilibrium Score Regression (LDSC) and Mendelian Randomization (MR) to assess the genetic correlation and causality between traits related to cardiac structure and sarcopenia, with further validation of the results using validation set data. In the study, we established a scoring system to identify high-confidence trait pairs. For these pairs, we conducted sensitivity analyses to assess their heterogeneity and pleiotropy. Additionally, we undertook follow-up studies of these trait pairs, including using mediation analysis to evaluate the potential mediating effects of lipids, and enrichment analysis to explore possible shared biological pathways linking these characteristics.

**Results:** The genetic correlation analyses during the discovery and validation phases identified 5 pairs (in forward analysis) and 16 pairs (in reverse analysis) of high-confidence trait pairs. As a key sarcopenia-related feature, appendicular lean mass (ALM) exhibited positive causal relationships with several cardiac structural features, including the volumes of the left and right atria. Mediation analysis suggested that certain lipids, such as phosphatidylcholine, might mediate these causal relationships. Notably, gene and pathway enrichment analyses revealed that genes associated with significant SNPs in high-confidence trait pairs were enriched in multiple key biological processes and pathways, such as tube morphogenesis, cardiac right ventricle morphogenesis, and muscle system processes in the forward analysis, as well as skeletal system development, heart development, and hemopoiesis in the reverse analysis. Cell-type enrichment analysis pointed to endothelial cells, smooth muscle cells, fibroblasts, and mesenchymal stem cells.

**Conclusions:** This study provides evidence for genetic relationships between cardiac structures and sarcopenia-related traits. These findings may enhance our understanding of the biological mechanisms underlying age-related diseases and provide scientific basis for developing future preventive and therapeutic strategies.

## Introduction

As global population aging intensifies, there has been a notable increase in the incidence of age-related diseases, particularly sarcopenia and cardiovascular diseases (CVDs) (1). These conditions, common in elderly syndromes, are characterized by high incidence rates, subtle onset, and wide-ranging impacts on the body, leading to significant burdens on both family healthcare and public health expenditures (2-4). Sarcopenia is defined as the age-associated decline in skeletal muscle mass, strength, or physical performance (5). This condition results in frailty, impaired mobility, diminished quality of life, and increased mortality risk. The development of sarcopenia is linked to several factors including chronic inflammation, hormonal fluctuations, malnutrition, and decreased physical activity (6). Additionally, the association between sarcopenia and various chronic diseases, especially CVDs, has been extensively studied and acknowledged. CVDs rank among the top causes of death worldwide. From 1990 to 2019, the number of people affected by CVDs rose from 271 million to 523 million, with associated deaths increasing from 12.1 million to 18.6 million (7).

In recent years, the link between CVDs and sarcopenia has garnered increasing attention from researchers. A meta-analysis incorporating 22 articles and covering 4,327 patients revealed that the prevalence of sarcopenia among individuals with CVDs varied from 10.1% to 68.9%, with an overall pooled prevalence of 35%. In comparison, the prevalence in the general population ranged from 2.9% to 28.6%, with a pooled prevalence of 13%. These findings indicate that the prevalence of sarcopenia is approximately twice as high in CVD patients as in the general population, suggesting a positive correlation between CVDs and sarcopenia (8). Additionally, a longitudinal study of 15,000 middle-aged and elderly Chinese individuals (aged≥ 45 years) showed that compared to their healthy counterparts, those with sarcopenia and possible sarcopenia had a 72% and 29% increased incidence of CVDs, respectively. The risk of new cardiovascular events was also higher, increasing by 33% and 22% respectively (9). The intricate relationship between sarcopenia and CVDs may involve multiple pathophysiological mechanisms, including inflammation, insulin resistance, oxidative stress, endothelial dysfunction, and hormonal changes (1).

Research has demonstrated that alterations in cardiac structure and function in the elderly are closely linked to sarcopenia. Factors such as lipid metabolism, inflammatory responses, and oxidative stress may play significant roles in the pathogenesis linking these cardiac changes to sarcopenia, prompting an exploration of the cardiac-skeletal muscle axis (10, 11). Studies have shown a significant positive correlation between left ventricular mass and limb skeletal muscle mass in elderly populations exhibiting reduced muscle mass and physical function (12). Furthermore, a large-scale study involving 67,106 young and middle-aged Korean individuals revealed a strong, independent link between low relative skeletal muscle mass and an increased incidence of left ventricular diastolic dysfunction (13). Another investigation found that in elderly individuals with preserved cardiac function, sarcopenia was associated with decreased volumes of both the left ventricle and the left atrium (14). Despite epidemiological studies showing a link between cardiac structural abnormalities and sarcopenia, the causal relationships and underlying biological mechanisms between these conditions are not yet fully understood. Traditional observational studies, constrained by confounding factors, struggle to resolve these complexities.

In recent years, genetic correlation analysis and MR have emerged as pivotal methods for investigating potential causal relationships among genetic phenotypes. MR, which bridges the gap between traditional epidemiological studies and randomized controlled trials, offers an intermediate level of evidence. Compared to case-control or cohort studies, MR provides more robust evidence (15). By employing genetic variants as instrumental variables, MR effectively minimizes the confounding factors commonly encountered in traditional observational studies, thereby enabling the assessment of potential causal effects from one phenotype to another (16). Utilizing genetic epidemiological models, MR can accurately estimate the association strength between exposures and outcomes, producing results that are less prone to confounding and avoiding reverse causation issues. Another key tool in genetic correlation analysis is LDSC. LDSC differentiates the statistical inflation caused by polygenic foundations from associations due to population stratification or other confounding factors. It is used to estimate heritability and genetic correlations between traits based on Single Nucleotide Polymorphism (SNP) data (17). This method provides a robust tool for dissecting the genetic architecture and interactions of complex traits.

This study utilized MR analysis to investigate the potential reciprocal causal relationships between cardiac structural phenotypes and sarcopenia-related traits, aiming to uncover any possible bidirectional causality between these two conditions. Additionally, the study included cross-trait genetic correlation analysis to estimate the genetic correlations between cardiac structures and sarcopenia-related traits, thereby further exploring their shared genetic bases. Through the above analysis, we determined the high confidence trait pairs. Furthermore, considering the potential mediating roles of biomarkers in pathological processes, this research carried out mediation analysis on 179 plasma lipid species to determine their involvement in the causal pathways between cardiac structures and sarcopenia traits. To elucidate the shared biological pathways underlying these cardiometabolic-musculoskeletal interactions, enrichment analysis is integrated to identify tissue-specific regulatory networks and functionally annotated gene sets. The objective of this research is to provide a more scientific foundation for future preventative and therapeutic strategies targeting these pathological conditions and to offer new insights into the biological bases and common pathological mechanisms of these prevalent diseases in the elderly. The findings of this study are expected to advance further research in the domains of cardiac and muscular health in the elderly, ultimately enhancing their health and quality of life.

## Methods

### 1. Study Design

In this study, first, we used the cross-trait LDSC method to estimate the genetic correlation between cardiac structure and sarcopenia. Discovery and confirmatory sets are set for all sarcopenia-related traits. Then two-sample MR analysis was crafted in accordance with the STROBE-MR guidelines (Supplementary STROBE checklist) and used to evaluate the causal relationship between exposures and outcomes. In order to obtain impartial results, three hypotheses of MR must be met: (a) the genetic instrumental variables (IVs) should be highly correlated with exposure; (b) there is no confounding of the instrument-outcome association; (c) the genetic IVs affected the outcome only through exposure but not through other biological pathways (18). Therefore, the genome-wide significance single nucleotide polymorphisms (SNPs, p ≤5×10^-8^) were used as IVs, and then a series of analyses were carried out, including four MR methods (inverse-variance weighted (IVW), simple mode, weighted median and weighted mode), as well as heterogeneity and horizontal pleiotropy tests (such as Cochran’s Q test, MR-Egger intercept test and MR-PRESSO test) to verify our results. Bidirectional MR analysis was performed to estimate the potential reverse causal impact of sarcopenia on heart structure. Finally, we conducted follow-up analysis, including mediation analysis and enrichment analysis, to explore the possible mechanisms of action between traits. The whole workflow of MR analysis was shown in **Figure 1**.

**Figure 1.**
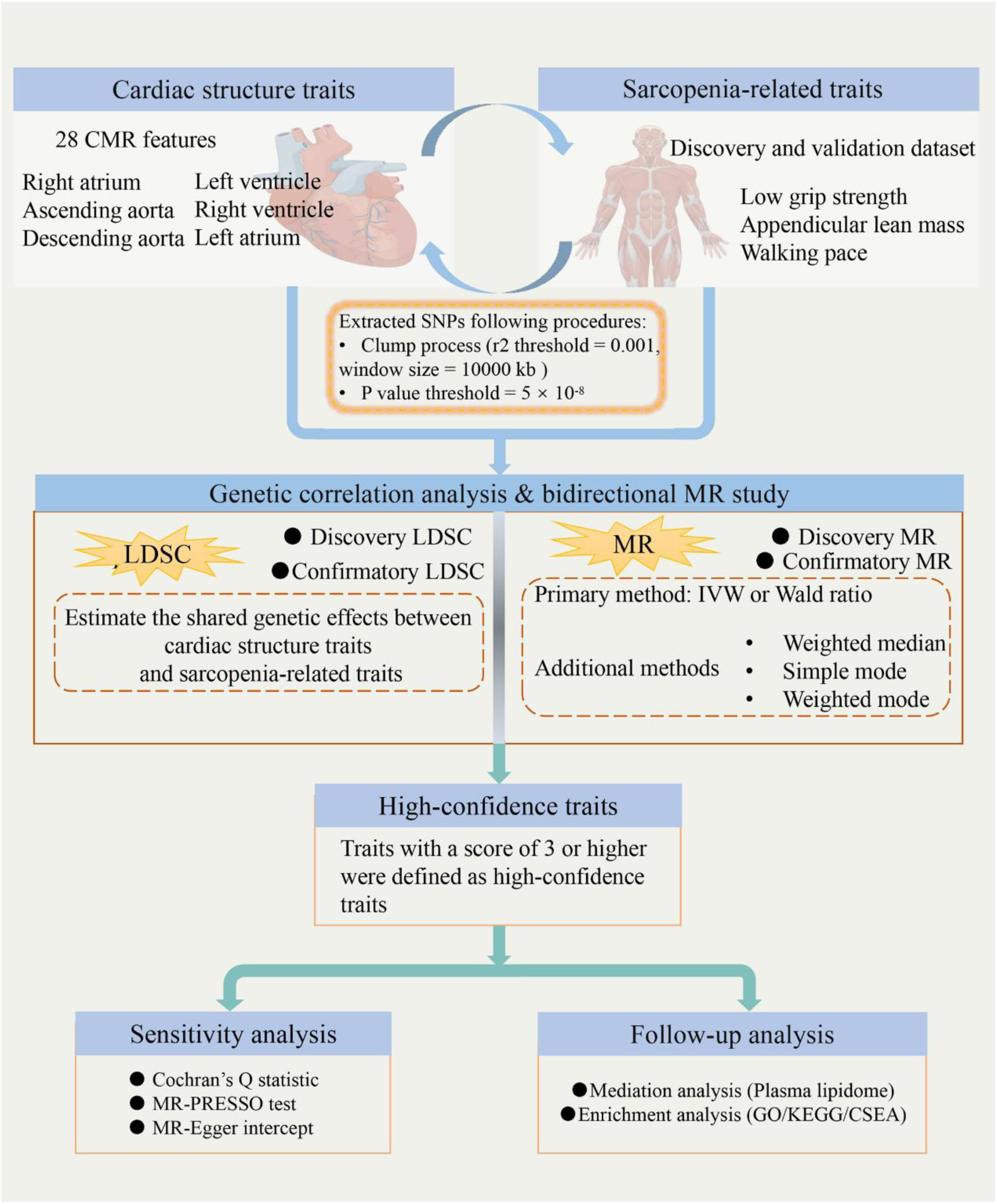
Overview of analyses performed in this study.

### 2. Data Sources

All analyses were based on publicly shared databases and no additional ethical approvals were required.

#### 2.1 Cardiac magnetic resonance imaging data

Cardiac structural data were derived from 31,000 participants in the UK Biobank cohort (19). Cardiac magnetic resonance imaging (CMR) features were segmented and extracted, encompassing measurements from four cardiac regions—left ventricle (LV), right ventricle (RV), left atrium (LA), and right atrium (RA)—as well as two segments of the aorta: the ascending aorta (AAo) and descending aorta (DAo). Additionally, regional phenotypes of left ventricular strain were included (20, 21). The GWAS results for 28 CMR traits are accessible via http://heartkp.org/.

#### 2.2 Sarcopenia-related traits data

Summary data on low grip strength (LGS) were sourced from a meta-analysis conducted by Garan Jones (22), comprising 256,523 individuals of European ancestry aged 60 years and older. Discovery dataset adhered to the stringent Foundation for the National Institutes of Health (FNIH) criteria (men <26 kg, women <16 kg). Validation employed the European Working Group on Sarcopenia in Older People (EWGSOP) criteria (grip strength <30 kg for men, <20 kg for women). Among 48,596 cases identified by the EWGSOP criteria, 20,335 met the stricter FNIH standard. Lean body mass, a key metric of muscle quality in older adults and extensively studied in sarcopenia (23), was evaluated using appendicular lean mass (ALM). GWAS summary statistics for ALM were obtained from the UK Biobank (n = 205,513) and further validated with an additional dataset of 244,730 individuals of European ancestry, adjusting for appendicular fat mass, age, and other covariates (24). Two GWAS summary data for walking pace (WP) were derived from Neale Lab (ukb-a-513) and the MRC-IEU Consortium (ukb-b-4711), comprising 335,349 and 459,915 individuals of European ancestry, respectively.

#### 2.3 Plasma lipidome data

To minimize bias, lipidomic data from an in-depth analysis of 179 plasma lipidome in a Finnish cohort of 7174 from sources different from the exposure and outcomes database were selected (25). We performed a mediation analysis of these 179 liposomes. Features are accessible online at https://gwas.mrcieu.ac.uk/ (accession numbers: GCST90277238 to GCST90277416).

### 3. LDSC

Genetic correlation analyses were performed to estimate the shared genetic effects between traits and to highlight underlying common etiologies. LDSC provided a robust method to evaluate heritability and genetic correlations based on GWAS summary data while distinguishing true polygenicity from confounding biases such as population stratification and cryptic relatedness. Using LDSC software (https://github.com/bulik/ldsc), we conducted cross-trait LDSC to estimate the genetic correlation (rg) between cardiac structure and sarcopenia. The genetic correlation (ranging from −1 to 1) between two traits was calculated using the known linkage disequilibrium (LD) structure of European ancestry reference data from the 1000 Genomes Project (26).

### 4. Genetic Instruments Selection

The screening of SNPs was conducted following the procedures outlined as follows: (1) To ensure statistical significance, a genome-wide threshold of p≤ 5×10^-8^ was applied, due to the limited number of SNPs screened when selecting this threshold for liposome data, the threshold was relaxed to 1×10^-5^; (2) SNPs with minor allele frequencies < 0.05 were excluded from analysis; (3) SNPs featuring mismatched alleles, such as C/T and C/A, were excluded; (4) Palindromic variants with equivocal strands, such as C/G or A/T, were also removed from the dataset; (5) The genetic variations that passed the aforementioned filters were further processed in Plink to identify lead SNPs using specific parameters for clumping: window size = 10000 kb and r^2^ < 0.001.

### 5. MR Analysis

Based on the results of LDSC, we employed bidirectional MR to further investigate the genetic causal relationship between cardiac structure and sarcopenia. The study was divided into discovery and validation phases. SNPs with low F-statistics (<10) were excluded from our analysis to avoid significant weak instrument bias. When only a single SNP was available, the Wald Ratio (WR) method was applied, whereas for two or more SNPs, the multiplicative random effects IVW approach was utilized as the primary method for estimating the causal relationship between exposure and outcome. This method is considered the most efficient analysis when valid instrumental variables are present, and the multiplicative random effects model outperforms additive random effects and fixed-effects models under conditions of excessive heterogeneity (27).

When the number of SNPs for a trait was three or more, three additional MR methods—simple mode, weighted median, and weighted mode—were employed for sensitivity analyses as supplements to the main analysis results. In this study, the results of causal relationships are primarily based on the results of the primary analysis, while the results of supplementary analysis require consistency with the primary analysis. For traits with high confidence, Cochran’s Q test was further conducted to examine the heterogeneity of IVW estimates. To evaluate the detectability of horizontal pleiotropy, we applied the MR-Egger method by assessing the intercept of the genetic instruments. The MR-PRESSO approach was implemented to detect and correct pleiotropy and potential outliers, including both global and outlier tests, and causal estimates were recalculated after removing identified outliers.

### 6. Scoring of potential effector traits

Combining the results of LDSC and MR, we assigned a simple scoring system to the positive traits identified in the study: traits deemed significant in either the discovery LDSC or MR analysis were scored 1 point; traits significant in any validation phase or in both discovery LDSC and MR analyses were scored 2 points; traits significant in both discovery LDSC and MR analyses, with at least one result validated, were scored 3 points; and traits significant in both discovery and validation phases of LDSC and MR were scored 4 points. Traits with a score of 3 or higher were defined as high-confidence traits.

### 7. Mediation analysis

Additionally, mediation MR analysis was conducted to explore whether 179 lipidomic traits mediated the causal pathways from exposure to outcomes for these high-confidence traits. This mediation analysis was performed in two steps: the first step assessed the causal association between the exposure and mediators, and in the second step, meaningful mediators were tested for their causal association with the outcome. Mediators demonstrating significance in both steps (p ≤ 0.05) were considered potential mediators. Then the mediation proportion and p-values were calculated, and mediators with p ≤ 0.05 were regarded as significantly mediating factors.

### 8. Enrichment analysis

In our study, we initially selected SNPs with significant associations in high-confidence trait pairs from genomic data, cataloging and annotating these SNPs using the public database dbSNP (https://www.ncbi.nlm.nih.gov/snp/). Subsequently, we conducted a comprehensive enrichment analysis of the functions and pathways of the corresponding genes, aiming to uncover the shared biological pathways and potential roles they play in disease mechanisms. Specifically, using Metascape (https://metascape.org)(28), we performed pathway and process enrichment analyses, including GO Biological Processes, KEGG Pathway, GO Molecular Functions, GO Cellular Components, Reactome Gene Sets, Hallmark Gene Sets, Canonical Pathways, BioCarta Gene Sets, CORUM, and WikiPathways. Using the online platform WebCSEA (Web-based Cell-type Specific Enrichment Analysis), we analyzed CSEA for 1,355 tissue cell types from 61 different general tissues across 11 human organ systems(29). These integrative methods not only enhanced our systemic understanding of gene or cell interactions but also facilitated the extraction of key biological insights from a complex genetic backdrop by elucidating the associations among gene functions, pathways, and cells.

### 9. Statistical analysis

The MR and additional analyses were conducted using the “Two Sample MR” package in R (version 4.3.1). Results were expressed as odds ratios (ORs) with their respective 95% confidence intervals (CIs) per standard deviation. The mediation proportion was derived using a formula where β represents the total effect obtained from the initial analysis, and β1 and β2 represent the effects of the exposure trait on the mediator and the mediator on the outcome, respectively. Thus, the formula is: Proportion = (β1 × β2) / β.

## Results

### 1. Genetic correlation

First, a discovery genetic correlation analysis was performed using GWAS summary data of ALM, WP and FNIH-defined LGS, applying stricter standards (**Figure 2** and **Supplemental table 1**). Among 28 CMR traits, we identified that 28% of the genetic components of CMR were associated with LGS, 64% with ALM, and 43% with WP (p ≤ 0.05). To enhance confidence in our findings, confirmatory LDSC analysis was conducted using LGS defined by the EWGSOP standard, along with alternative datasets for ALM and WP. LDSC identified genetic correlations between 2 (left ventricular stroke volume [LVSV], right atrium maximum volume [RAV_max]), 15 (excluding global myocardial-wall thickness at end-diastole [WT_global], descending aorta maximum area [DAo_max_area], descending aorta minimum area [DAo_min_area]), and 11 (excluding descending aorta distensibility [DAo_aortic_distensibility]) CMR traits and LGS, ALM, and WP from the discovery analysis. Across six datasets, LVSV and RAV_max consistently demonstrated significant genetic correlations with LGS, ALM, and WP.

**Figure 2.**
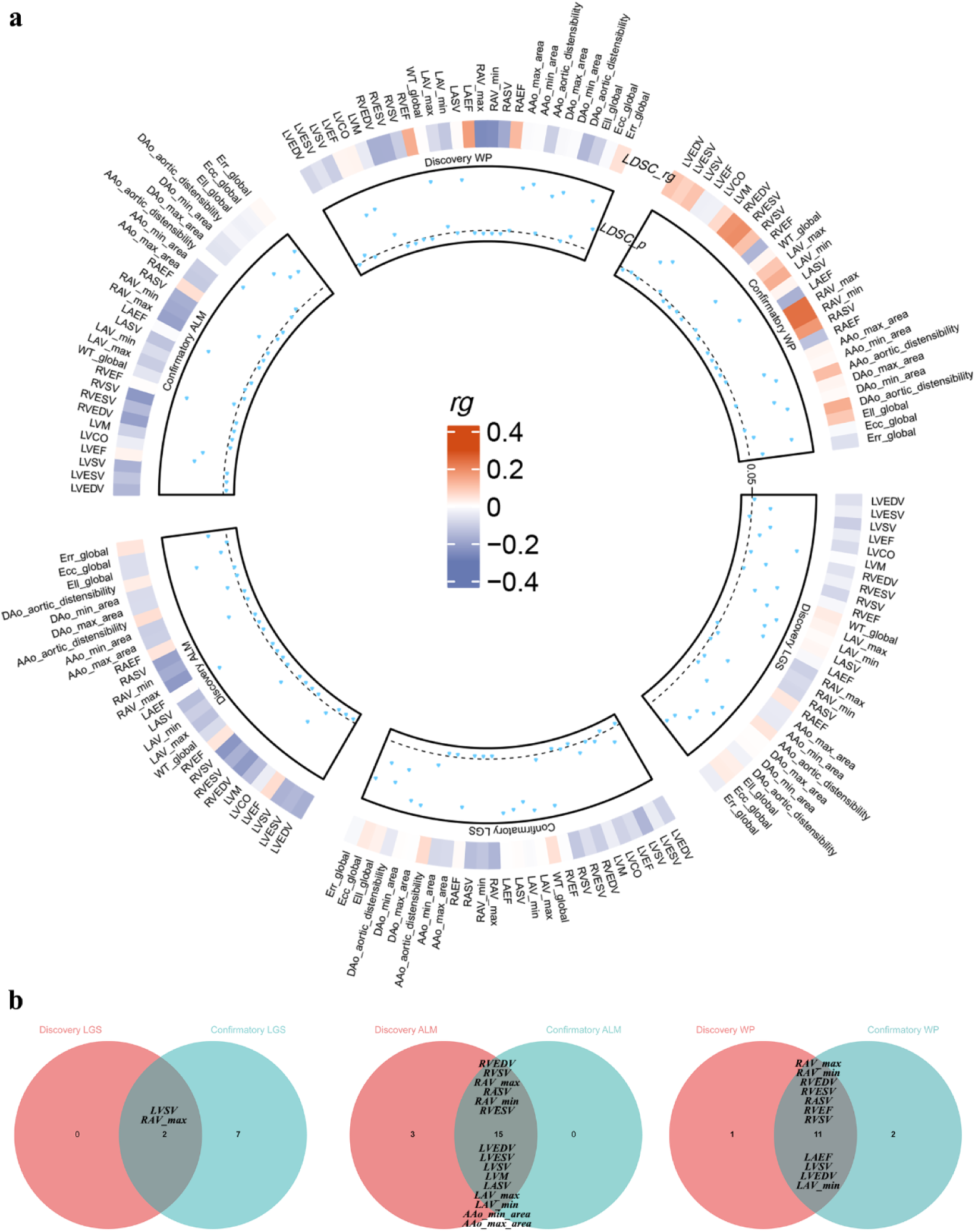
Global genetic correlation between cardiac structures and sarcopenia-related traits. **(a)** Circular heatmap and scatter plot of results from Linkage Disequilibrium Score Regression (LDSC) showing the global genetic correlation (rg) between 28 cardiac structures from cardiac magnetic resonance imaging and sarcopenia-related traits [low grip strength (LGS), appendicular lean mass (ALM), and walking pace (WP)]. **(b)** Venn diagram of the intersection results of discovery and confirmatory LGS, ALM and WP.

### 2. Influence of heart structure on sarcopenia-related traits

#### 2.1 Exploring the causal effects of heart structure on sarcopenia for novel discoveries

For LGS as defined by the FNIH standard, the primary MR results revealed that five CMR traits (RAV_max, right ventricular stroke volume [RVSV], left ventricular end-systolic volume [LVESV], WT_global, and ascending aorta maximum area [AAo_max_area]) exhibited negative associations with LGS. In sensitivity analyses, the Weighted Median method supported the association for RAV_max (p = 0.03), while the Simple Mode method supported the result for LVESV (p = 0.05). For ALM, the IVW model identified negative causal associations between three traits (left atrium minimum volume [LAV_min], ascending aorta minimum area [AAo_min_area], and AAo_max_area) and ALM, whereas global longitudinal strain (Ell_global) demonstrated a positive causal relationship with ALM. For traits involving multiple SNPs (AAo_min_area and AAo_max_area), the Simple Mode method corroborated the association for AAo_min_area (p = 0.03). Similarly, MR analysis identified causal relationships between five traits (AAo_max_area, AAo_min_area, Ell_global, RAV_max, and WT_global) and WP, with only RAV_max showing a positive association. The Weighted Median method also supported the results for AAo_max_area (p = 0.02) and RAV_max (p = 0.02). All other analytical methods demonstrated directions consistent with the IVW results (**Figure 3** and **Supplemental table 2**).

**Figure 3.**
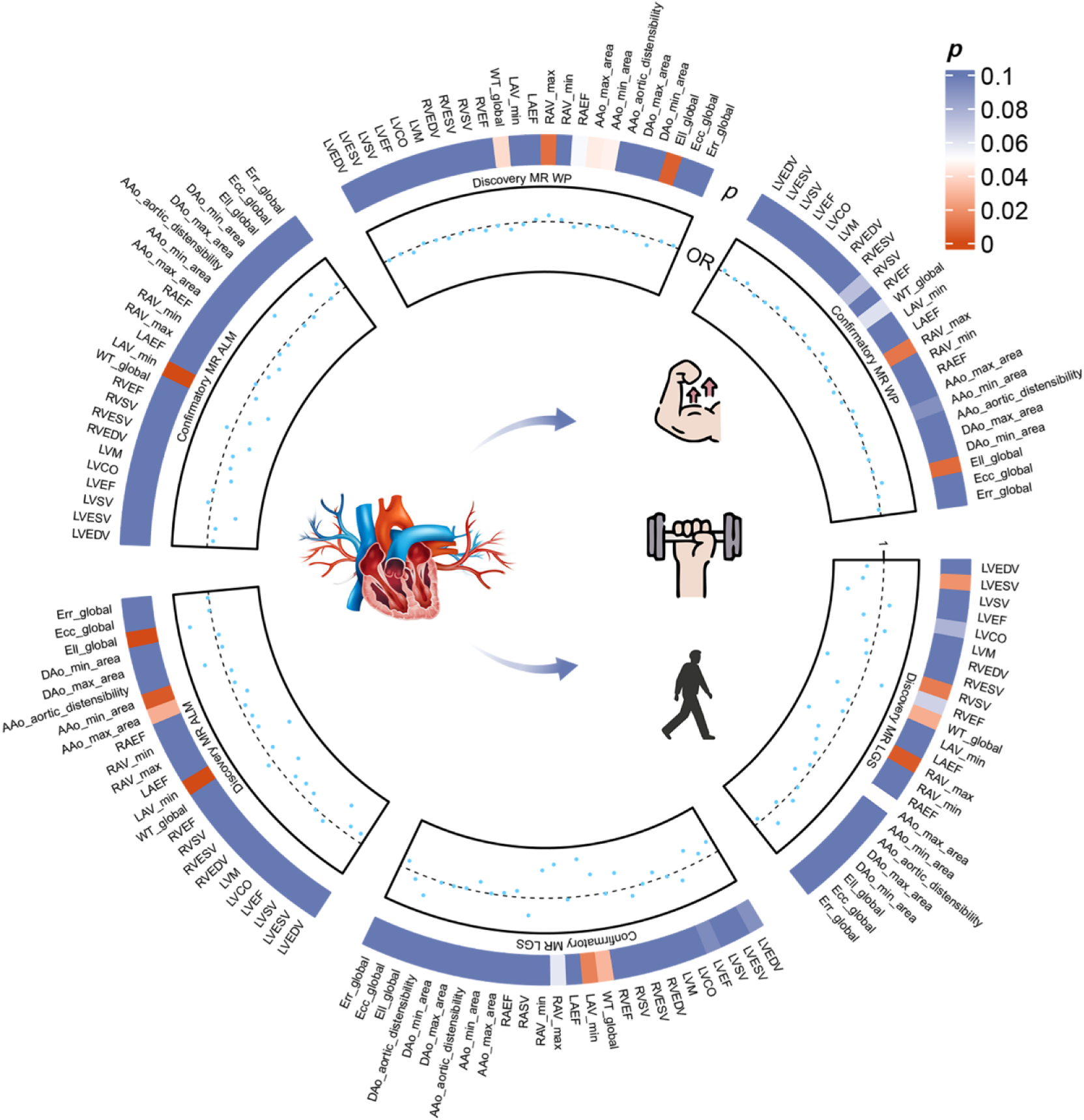
Effects of cardiac structures on sarcopenia-related traits. Circular heatmap and scatter plot of results from MR analyses showing the potential causal effects of cardiac structures on sarcopenia-related traits.

#### 2.2 Validating the causal effects of heart structure on sarcopenia

For LGS, datasets based on the EWGSOP standard, as well as alternative ALM and WP data, were utilized for confirmatory MR analysis. In the confirmatory MR, increased volumes of three cardiac structural traits were associated with lower grip strength, among which one trait (LVESV) was also identified as having a causal relationship in the discovery MR analysis. Only one cardiac structure (LAV_min) demonstrated a negative causal relationship with low grip strength, an association that was also confirmed in the discovery phase. Furthermore, two traits (Ell_global and RAV_max) identified in the discovery phase as being associated with WP were similarly validated (**Figure 3** and **Supplemental table 2**).

### 3. Influence of sarcopenia-related traits on heart structure

In reverse MR, analyses were also divided into discovery and validation phases (**Figure 4** and **Supplemental table 3**). Among the 28 CMR traits, the discovery phase revealed causal relationships with 17, 1, and 0 traits for ALM, LGS, and WP, respectively. In the ALM confirmatory dataset, 16 traits were found to have causal relationships, 15 of which (excluding left atrium maximum volume [LAV_max]) were also confirmed in the discovery phase. Results from both the discovery and validation phases consistently demonstrated positive causal associations between ALM and these 15 traits. However, the other two validation datasets failed to replicate the findings of the discovery phase.

**Figure 4.**
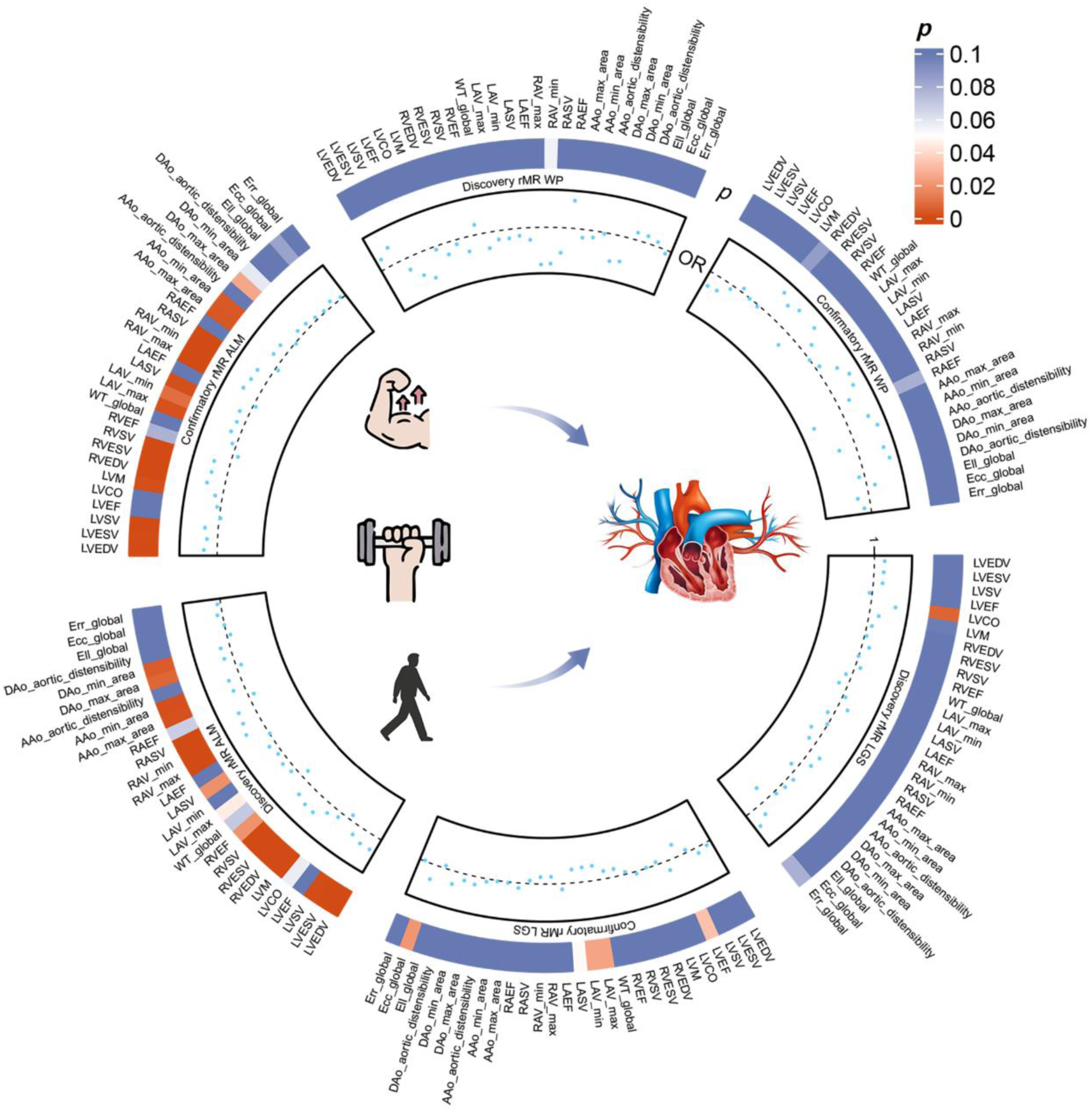
Effects of sarcopenia-related traits on cardiac structures. Circular heatmap and scatter plot of results from rMR analyses showing the potential causal effects of sarcopenia-related traits on cardiac structures.

### 4. Scoring for high confidence traits

Here, a new scoring system has been constructed. As shown in **Table 1** and **Figure 5**, combining results from the discovery and validation phases of LDSC and MR analyses allowed us to identify several high-confidence traits (traits with a score of 3 or higher). In the forward analysis, we identified three high-confidence CMR traits (AAo_max_area, AAo_min_area, and LAV_min) influencing ALM, as well as RAV_max influencing both LGS and WP. In reverse MR, the results demonstrated that ALM has causal relationships with 16 high-confidence CMR traits. **Figure 6** shows the interaction patterns between these CMR and the traits associated with sarcopenia.

**Figure 5.**
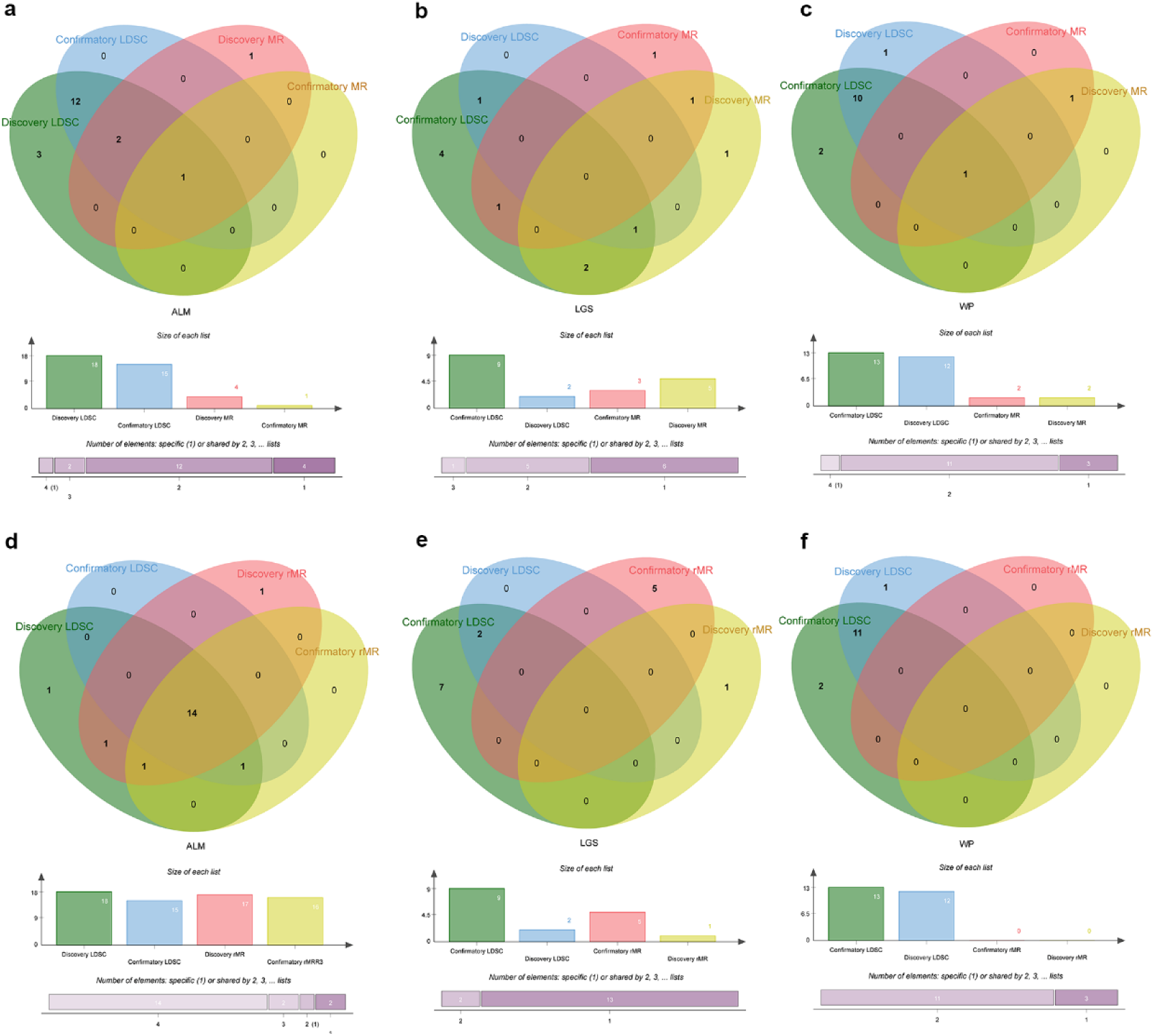
The Venn diagram of the discovery and confirmatory MR (a, b, c) and rMR(d, e, f) on ALM, LGS and WP. Low grip strength (LGS), appendicular lean mass (ALM), walking pace (WP)

**Figure 6.**
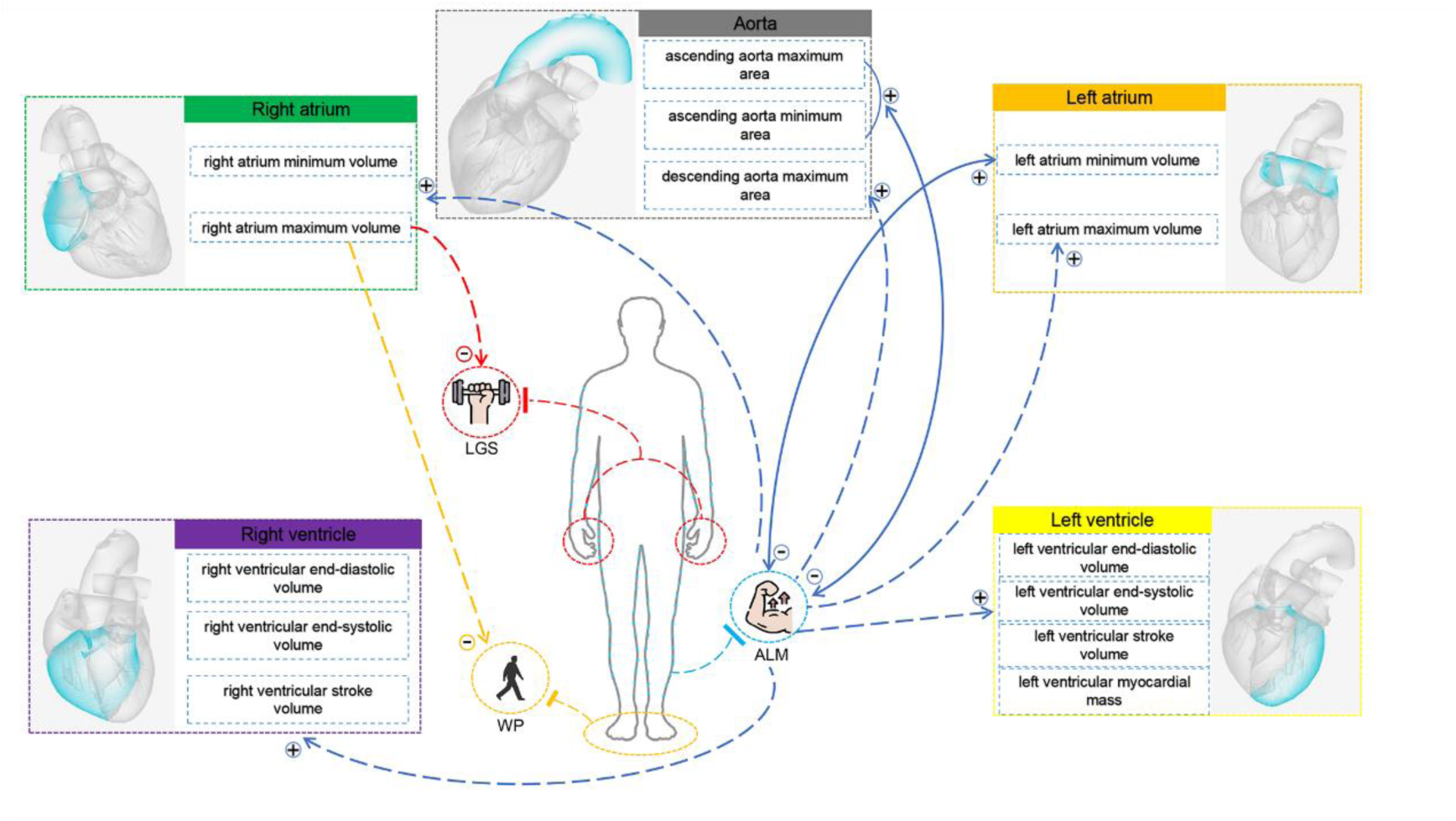
Interaction pattern between cardiac structures and sarcopenia-related traits. The solid line represents the bidirectional association. LGS, low grip strength; ALM, appendicular lean mass; WP, walking pace

**Table 1.** High credibility traits scoring table.

| Exposure\|Name |  | Outcome\|Name |  | LDSC |  | MR |  | Score |
| --- | --- | --- | --- | --- | --- | --- | --- | --- |
|  |  |  |  | Discovery | Confirmatory | Discovery | Confirmatory |  |
| Forward Mendelian randomization |  |  |  |  |  |  |  |  |
| ascending aorta maximum area | AAo_max_area | appendicular lean mass | ALM | ✓ | ✓ | ✓ | □ | 3 |
| ascending aorta minimum area | AAo_min_area | appendicular lean mass | ALM | ✓ | ✓ | ✓ | □ | 3 |
| right atrium maximum volume | RAV_max | low hand grip strength | LGS | ✓ | ✓ | ✓ | □ | 3 |
| left atrium minimum volume | LAV_min | appendicular lean mass | ALM | ✓ | ✓ | ✓ | ✓ | 4 |
| right atrium maximum volume | RAV_max | Usual walking pace | WP | ✓ | ✓ | ✓ | ✓ | 4 |
| Reverse Mendelian randomization |  |  |  |  |  |  |  |  |
| appendicular lean mass | ALM | descending aorta maximum area | DAo_max_area | ✓ | ✓ | ✓ | □ | 3 |
| appendicular lean mass | ALM | left atrium minimum volume | LAV_min | ✓ | ✓ | □ | ✓ | 3 |
| appendicular lean mass | ALM | left ventricular end-diastolic volume | LVEDV | ✓ | ✓ | ✓ | ✓ | 4 |
| appendicular lean mass | ALM | left ventricular end-systolic volume | LVESV | ✓ | ✓ | ✓ | ✓ | 4 |
| appendicular lean mass | ALM | left ventricular stroke volume | LVSV | ✓ | ✓ | ✓ | ✓ | 4 |
| appendicular lean mass | ALM | left ventricular myocardial mass | LVM | ✓ | ✓ | ✓ | ✓ | 4 |
| appendicular lean mass | ALM | right ventricular end-diastolic volume | RVEDV | ✓ | ✓ | ✓ | ✓ | 4 |
| appendicular lean mass | ALM | right ventricular end-systolic<br>volume | RVESV | ✓ | ✓ | ✓ | ✓ | 4 |
| appendicular lean mass | ALM | right ventricular stroke volume | RVSV | ✓ | ✓ | ✓ | ✓ | 4 |
| appendicular lean mass | ALM | left atrium maximum volume | LAV_max | ✓ | ✓ | ✓ | ✓ | 4 |
| appendicular lean mass | ALM | left atrium stroke volume | LASV | ✓ | ✓ | ✓ | ✓ | 4 |
| appendicular lean mass | ALM | right atrium maximum volume | RAV_max | ✓ | ✓ | ✓ | ✓ | 4 |
| appendicular lean mass | ALM | right atrium minimum volume | RAV_min | ✓ | ✓ | ✓ | ✓ | 4 |
| appendicular lean mass | ALM | right atrium stroke volume | RASV | ✓ | ✓ | ✓ | ✓ | 4 |
| appendicular lean mass | ALM | ascending aorta maximum area | AAo_max_area | ✓ | ✓ | ✓ | ✓ | 4 |
| appendicular lean mass | ALM | ascending aorta minimum area | AAo_min_area | ✓ | ✓ | ✓ | ✓ | 4 |

### 5. Heterogeneity and horizontal pleiotropy detection

For these high-confidence traits, we conducted heterogeneity and horizontal pleiotropy tests (**Supplemental table 4**). According to MR-Egger regression intercept analysis, there was no significant directional horizontal pleiotropy (p > 0.05). Cochran’s IVW Q-test indicated that most of the IVs for these associations exhibited heterogeneity (p ≤ 0.05), and the MR-PRESSO global test also detected outliers (RSSobs p ≤ 0.05). However, these outliers did not alter the directionality of the results (distortion test p > 0.05). Post-correction analyses confirmed that all associations remained significant (outliers corrected p ≤ 0.05).

### 6. Mediation analysis

#### 6.1 First step

Both forward and reverse mediation analyses were performed for all high-confidence traits to evaluate potential mediation effects of liposomes in the causal relationships between exposures and outcomes (**Supplemental table 5**). In the first-step analysis, we found that LAV_min, RAV_max, AAo_max_area, and AAo_min_area exhibited causal relationships with 9, 2, 4, and 4 liposomes, respectively, in forward analysis. Additionally, ALM from the discovery and confirmatory datasets showed causal relationships with 2 and 13 liposomes, respectively, in reverse analysis.

#### 6.2 Second step

For liposomes with causal effects, we further assessed their causal relationships with corresponding outcomes. The results (**Supplemental table 6**) revealed that for LAV_min, the lipid species Phosphatidylcholine (18:0_20:4) was causally associated with ALM in the confirmatory dataset, while Phosphatidylcholine (16:0_20:4) was causally associated with ALM in both the discovery and confirmatory sets. For AAo_max_area and AAo_min_area, Phosphatidylcholine (O-16:0_20:4) was causally associated with ALM in the discovery set. In the confirmatory ALM set, Sterol ester (27:1/16:0) was causally associated with LASV, Phosphatidylcholine (14:0_18:2) with RVESV, Phosphatidylcholine (18:1_0:0) with RVSV, and Cholesterol levels with both LVEDV and LVSV.

#### 6.3 Mediation proportion

The plots in **Figure 7** visually depict these liposomes with potential mediatory significance. Subsequently, we calculated the mediation proportions and p-values for these liposomes (**Supplemental table 7**). The results indicated that these plasma lipidomes mediated 2.79%–5.68% of the causal effects. However, none of which demonstrated statistically significant mediation effects (p > 0.05).

**Figure 7.**
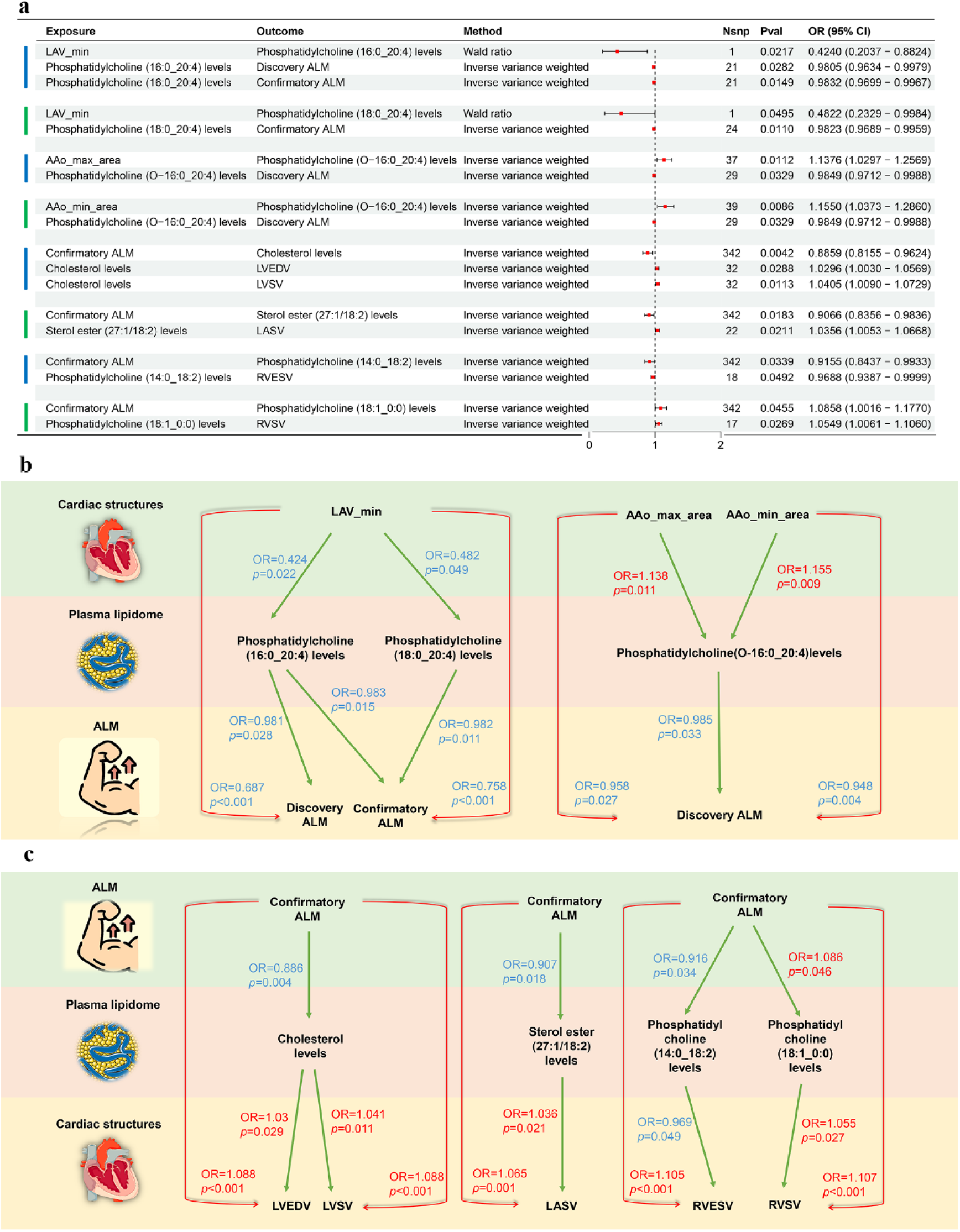
Mediation analyses show causal effects of 8 liposomes on ALM and cardiac structures. OR values indicate the causal effect estimates using the inverse variance weighted method (truncated at p<0.05).

### 7. Enrichment analysis

SNPs significantly associated in both forward and reverse analyses within high-confidence trait pairs were translated into genes and underwent enrichment analysis, as shown in **Supplemental Table 8**. In the forward analysis (**Supplemental Figures 1 and 2**), the top six enriched clusters included tube morphogenesis, cardiac right ventricle morphogenesis, muscle system process, negative regulation of muscle cell apoptotic process, sensory organ morphogenesis and localization within membrane (AAo_max_area→ALM); angiogenesis, PID AP1 PATHWAY, muscle contraction, extracellular matrix, phospholipase C-activating G protein-coupled receptor signaling pathway and localization within membrane (AAo_min_area→ALM).

In the reverse analysis (**Supplemental Figures 3 and 4**), the top six enriched clusters included Cancer pathways, skeletal system development, small cell lung cancer, hemopoiesis, Epstein-Barr virus infection and endochondral ossification (discovery ALM→cardiac structures); Pathways in cancer, skeletal system development, transcription regulator complex, heart development, epithelial cell differentiation and transcription factor binding. (confirmatory ALM→cardiac structures), detailed in **Supplemental Table 9**.

CSEA indicated that in the forward analysis, these genes were predominantly enriched in hepatic stellate cells, endothelial cells, and smooth muscle cells (common cell types in the top 5, **Supplemental Figures 5 and 6**). In the reverse analysis, these genes were mainly enriched in fibroblast cells, mesenchymal stem cells, and endothelial cells (**Supplemental Figures 7 and 8**).

## Discussion

This study utilized LDSC and MR analysis to explore the genetic links between cardiac structural features and sarcopenia-related traits (GS, ALM, and WP). Through a scoring system, the research confirmed reliable significant genetic associations between cardiac structural features and sarcopenia-related traits. Several cardiac features, such as maximum right atrial volume and left ventricular end-systolic volume, demonstrated potential positive causal effects on GS and WP. Moreover, reverse MR analysis suggested that ALM has a positive causal influence on multiple cardiac structural features. Additionally, the study investigated the mediating role of lipid metabolism within these causal relationships, some plasma lipids, such as phosphatidylcholine, were identified as potential mediators. Enrichment analysis further provided potential insights into the biological pathways and processes underlying these relationships. Gene and pathway enrichment analyses revealed that genes associated with significant SNPs in high-confidence trait pairs were enriched in multiple key biological processes and pathways, such as tube morphogenesis, cardiac right ventricle morphogenesis, and muscle system processes in the forward analysis, as well as skeletal system development, heart development, and hemopoiesis in the reverse analysis. CSEA pointed to endothelial cells, smooth muscle cells, fibroblasts, and mesenchymal stem cells.

This study has revealed significant genetic correlations and causal relationships between various cardiac structural features and sarcopenia-related traits. Specifically, a notable positive correlation was observed between left ventricular end-systolic volume (LVESV) and both GS and ALM. Furthermore, RAV_max also showed positive correlations with GS and WP. These findings suggest that optimizing cardiac structures could have a direct impact on muscle health and overall physical performance. The left ventricle, as the heart’s primary pumping chamber, plays a crucial role in systemic circulation and oxygen delivery. Enhanced function of the left ventricle typically suggests improved blood circulation and oxygen supply, which are essential for maintaining and enhancing muscle function (30, 31). Moreover, an increase in cardiac output may aid in muscle mass development by improving metabolic efficiency and facilitating muscle repair (32-34). Therefore, an increase in LVESV may positively influence muscle function by enhancing systemic circulation and muscular oxygenation. Our results are supported by previous studies on cardiac structures and muscle functions. A sports cardiology screening program in Hungary, which included 425 healthy elite athletes (304 of whom were national team members in their age groups) and 55 age- and sex-matched healthy non-athletes as controls, found significant differences in left and right ventricular morphological and functional parameters between the two groups. Athletes exhibited notably higher indices of diastolic and systolic volumes (35). Another study utilizing the UK Biobank reported that higher levels of GS were associated with larger left ventricular end-diastolic volumes. However, after adjusting for all covariates, there was no statistically significant association between baseline GS and LVESV (36). This discrepancy could be attributed to racial differences, and our study, with its analysis involving a larger number of SNPs, possesses greater statistical power to explore these associations. The enlargement of RAV_max, typically linked with increased cardiac load, particularly under conditions of elevated right ventricular preload, may reflect the body’s adaptive response to endurance demands and muscle strength loss. The increase in right atrial volume might be an attempt by the body to compensate for declining muscle function by boosting cardiac output. Endurance athletes have been reported to have significantly higher right heart measurements compared to matched controls, both strength athletes and healthy non-athletes (37). In resting conditions, indexed maximum volumes of both right and left atria in athletes are greater than in non-athletes (38), suggesting that an increase in right atrial volume in athletes may be considered a physiological adaptation.

Considering the potential reciprocal impacts of sarcopenia-related traits on cardiac structures, this study demonstrates significant positive causal relationships between ALM and various cardiac structural features. This implies that a decrease in muscle mass may not merely result from cardiac abnormalities but could also play a role in exacerbating them. Recent studies emphasize that higher ALM is linked to a reduced risk of coronary artery disease, underscoring its protective function in cardiovascular health (24, 39, 40). A national cohort study involving more than 3.7 million young adults indicated that an increase in predicted ALM or a decrease in predicted fat mass is associated with a lower risk of developing CVD. Conversely, a decrease in predicted ALM or an increase in predicted fat mass is linked to a higher risk of CVD (41). The decline in muscle mass might influence cardiac structure and function through various mechanisms, such as altering metabolic states, hormonal levels, and inflammatory status, thereby increasing the cardiac workload and potentially triggering or worsening adverse cardiac structural changes. Thus, maintaining adequate muscle mass is critical for cardiac health, especially in preventing cardiovascular diseases.

In the mediation analysis, we observed that specific plasma lipids, such as phosphatidylcholine, might serve as intermediaries between cardiac structural characteristics and sarcopenia traits. As a vital component of cell membranes and a key mediator in inflammation regulation and cell signaling, phosphatidylcholine highlights that the modulation of lipid metabolism could be a crucial mechanism linking cardiac structures and sarcopenia traits (42). Foods rich in phosphatidylcholine, glycerophosphocholine, choline, and L-carnitine are metabolized by the gut microbiota to produce trimethylamine (TMA), which is further oxidized in the liver to trimethylamine N-oxide (TMAO) by flavin-containing monooxygenase enzymes, affecting the cardiovascular system (43). Research indicates that three gut microbial metabolites of phosphatidylcholine—choline, TMAO, and betaine—can predict cardiovascular disease risk, with elevated plasma levels of TMAO being independently associated with cardiovascular events (44, 45). The production of TMAO by gut microbiota, through the metabolism of phosphatidylcholine, directly causes increased platelet reactivity and enhanced thrombus formation, thus elevating the risk of thrombotic diseases such as myocardial infarction and stroke (46). These findings underscore a significant link between phosphatidylcholine and cardiovascular disease development. Recent MR studies have also identified a close association between sarcopenia and 27 lipid metabolites, including phosphatidylcholine, with these lipids showing a positive causal relationship with muscle mass and strength (47, 48). Given phosphatidylcholine’s role in maintaining cell membrane integrity and function, theoretically, supplementation could benefit the muscle mass and function decline associated with sarcopenia. However, the potential negative impacts on cardiovascular health must be carefully considered when evaluating phosphatidylcholine supplementation as a treatment strategy for sarcopenia. Although initial mediation analyses suggested a causal connection between plasma lipids with cardiac structure and sarcopenia traits, further mediation proportion analyses did not reveal significant mediating effects of these lipids. This indicates that the interactions between cardiac structures and sarcopenia likely involve more complex biological pathways, with lipid metabolism being only one aspect. Future studies are warranted to delineate the precise roles and underlying mechanisms of lipids within this context.

This study delved into the molecular mechanisms of the interaction between cardiac structure and sarcopenia through enrichment analysis. In the forward analysis, genes related to AAo_max_area→ALM were found to be mainly enriched in cardiovascular development associated pathways, such as cardiac tube morphogenesis and right ventricle morphogenesis, suggesting that cardiac structural abnormalities might affect skeletal muscle blood supply and function by influencing cardiovascular development (49, 50). Also, these genes were associated with muscle system processes and pathways of inhibiting myocyte apoptosis, indicating that cardiac chamber formation abnormalities might exacerbate muscle atrophy through the release of pro - apoptotic factors (51). Meanwhile, genes related to AAo_min_area→ALM were found to be mainly enriched in muscle contraction and G protein-coupled receptor signaling pathways activated by phospholipase C, further underscoring the role of vascular function and signaling conduction in cardiac-muscle crosstalk (52). In the reverse analysis, genes related to ALM and cardiac structures were found to be significantly enriched in skeletal system development, heart development and hemopoiesis pathways, thereby affecting cardiac structure. Reduced muscle mass might lead to myocardial fibrosis through chronic inflammation. Meanwhile, myokines secreted by muscles might regulate the differentiation of cardiac mesenchymal stem cells via BMP/Wnt signaling, thereby influencing cardiac structure (53). Cell-type-specific enrichment analysis showed that in the forward analysis, the related genes were mainly enriched in hepatic stellate cells, endothelial cells, and smooth muscle cells. In the reverse analysis, they were mainly enriched in fibroblasts, mesenchymal stem cells, and endothelial cells. These findings construct a multi-dimensional regulatory framework for heart-skeletal muscle interaction, providing a theoretical basis for developing combined therapies targeting both heart failure and sarcopenia.

This study presents several limitations. Firstly, the datasets for sarcopenia and cardiac structure used in the MR and LDSC analyses originate from GWAS studies conducted on European populations. While this approach minimizes bias from population stratification, it simultaneously constrains the generalizability of our findings. Notably, genetic structures and the genetic underpinnings or risk loci of diseases may exhibit heterogeneity across different populations, such as those of Asian or African descent, compared to Caucasian populations. Secondly, in the MR analysis, genetic instrumental variables are employed to represent exposure factors. Given the constraints of the sample size and additive regression model used in this study, only a small proportion of the variance in exposure factors is explained. Consequently, detecting subtle causal effects between complex traits remains challenging. Lastly, the effect sizes derived from the genetic correlation analysis are merely estimates based on the current dataset and models. These should not be considered equivalent to or substitutes for effect sizes obtained from observational clinical studies. To derive the most clinically relevant evidence, it is crucial to integrate genetic correlation analysis with a variety of evidence sources, including classical epidemiological studies, real-world research, bibliometric reviews, and meta-analyses.

## Conclusion

The findings from this study using bidirectional MR and genetic correlation analysis reveal a significant genetic linkage between cardiac structure and sarcopenia-related traits, illustrating a bidirectional relationship. These results underscore the importance of the cardiac-skeletal muscle axis in heart health and muscle function interplay. We identified potential mediation effects of plasma lipids and explored the shared biological pathways through enrichment analysis, which enhances our understanding of the interactions between cardiac and muscular health. Further research is needed to clarify these relationships’ biological mechanisms and confirm their relevance across different populations, advancing knowledge in age-related health issues.

## List of abbreviations

MR: Mendelian randomization. GWAS: genome-wide association study. IVW: inverse-variance weighted. SNPs: single nucleotide polymorphisms. IVs: instrumental variables. LDSC: linkage disequilibrium score regression. ALM: appendicular lean mass. CVDs: cardiovascular diseases. CMR: cardiac magnetic resonance imaging. LV: left ventricle. RV: right ventricle. LA: left atrium. RA: right atrium. AAo: ascending aorta. DAo: descending aorta. LGS: low grip strength. FNIH: Foundation for the National Institutes of Health. EWGSOP: European Working Group on Sarcopenia in Older People. WP: walking pace. WR: wald ratio. ORs: odds ratios. CIs: confidence intervals. LVESV: left ventricular end-systolic volume. TMA: trimethylamine. TMAO: trimethylamine N-oxide. CSEA: cell-type specific enrichment analysis.

## Declarations

### Ethics approval and consent to participate

Not applicable. Our analysis used publicly available genome-wide association study (GWAS) summary statistics. No new data were collected, and no new ethical approval was needed.

### Consent for publication

Not applicable.

### Availability of data and materials

The datasets used and analyzed during the current study are available from the corresponding author on reasonable request.

### Authors’ Contributions

WC and XWQ contributed equally to this work. WC, XWQ, LYS and ZSS designed the study. WC, XWQ and ZSS performed the analyses. WC, XWQ, ZY and ZSS analyzed the data. WC, XWQ, ZY, LZ, YG, ZD, LYS and ZSS interpreted the data, wrote the manuscript and revised the manuscript. All the authors approved of the final version of the manuscript.

## Acknowledgements

The authors thank the Home for Researchers editorial team (www.home-for-researchers.com) for improving the English language in this manuscript. The authors thank the vector diagrams provided by www.vecteezy.com and Figdraw.

